# Preclinical evaluation of RQ3013, a broad-spectrum mRNA vaccine against SARS-CoV-2 variants

**DOI:** 10.1101/2022.05.10.491301

**Authors:** Shudan Tan, Xue Hu, Yufeng Li, Zihan Wu, Jinghua Zhao, Guoliang Lu, Zhaoli Yu, Binhe Du, Yan Liu, Li Li, Yuchen Chen, Ye Li, Yanfeng Yao, Xiaoyu Zhang, Juhong Rao, Ge Gao, Yun Peng, Hang Liu, Zhiming Yuan, Jia Liu, Qianran Wang, Hengrui Hu, Xiaobo Gao, Hui Zhou, Hang Yu, Yingjie Xu, Wei Yu, Lin Feng, Manli Wang, Chao Shan, Jing Lu, Jinzhong Lin

**Author notes:** These authors contributed equally: Shudan Tan, Xue Hu, Yufeng Li, Zihan Wu, Jinghua Zhao, Guoliang Lu. Correspondence: Wei Yu; Lin Feng; Manli Wang; Chao Shan; Jing Lu; Jinzhong Lin.

## Abstract

The global emergence of SARS-CoV-2 variants has led to increasing breakthrough infections in vaccinated populations, calling for an urgent need to develop more effective and broad-spectrum vaccines to combat COVID-19. Here we report the preclinical development of RQ3013, an mRNA vaccine candidate intended to bring broad protection against SARS-CoV-2 variants of concern (VOCs). RQ3013, which contains pseudouridine-modified mRNAs formulated in lipid nanoparticles, encodes the spike(S) protein harboring a combination of mutations responsible for immune evasion of VOCs. Here we characterized the expressed S immunogen and evaluated the immunogenicity, efficacy, and safety of RQ3013 in various animal models. RQ3013 elicited robust immune responses in mice, hamsters, and nonhuman primates (NHP). It can induce high titers of antibodies with broad cross-neutralizing ability against the Wild-type, B.1.1.7, B.1.351, B.1.617.2, and the omicron B.1.1.529 variants. In mice and NHP, two doses of RQ3013 protected the upper and lower respiratory tract against infection by SARS-CoV-2 and its variants. We also proved the safety of RQ3013 in NHP models. Our results provided key support for the evaluation of RQ3013 in clinical trials.

## Introduction

In the early effort to combat the COVID-19 pandemic, mRNA vaccines with the lipid nanoparticle (LNP) delivery system have achieved unprecedented success. Two mRNA vaccines from Moderna (mRNA-1273) and Pfizer-BioNTech (BNT162b2), encoding the prefusion stabilized full-length spike (S) protein of SARS-CoV-2, are now being widely used^1, 2^, and several other mRNA vaccine candidates are in clinical trials. However, the effectiveness of the two mRNA vaccines is waning significantly as the virus continues to evolve, generating new variants. Studies have shown that sera from BNT162b2 or mRNA-1273 recipients exhibit more than a 30-fold reduction in neutralization titers against the latest Omicron variant compared to against Wild-type^3^.

Previously, we developed three mRNA vaccine candidates against the wild-type SARS-CoV-2 expressing the receptor-binding domain(RBD), S, and virus-like particles(VLP). Both the VLP and S immunogen elicited potent immune responses in mice^4^. We generated a pool of optimized mRNA sequences of assorted modifications from which the optimal one carrying the pseudouridine modification was identified. As the SARS-CoV-2 variants of concern (VOCs) kept posing new waves of global health threats, we began to focus on developing a broad-spectrum mRNA vaccine. One candidate, RQ3013, adapted from the previous S-based mRNA vaccine, showed encouraging results in the preclinical development. The mRNA sequence and modification of RQ3013 are inherited from its predecessor, but several changes have been made to the S immunogen. The furin cleavage site was mutated to prevent proteolysis of the S protein. The C-terminal eighteen residues were truncated for enhanced antigen expression. Most importantly, the S antigen carries mutations from the B.1.1.7 and B.1.351 variants and shares seven of them with the S protein from B.1.1.529. Here we characterized the structure of expressed immunogen of RQ3013 and reported a comprehensive evaluation of the immunogenicity, efficacy, and safety of RQ3013 in animal models of mice, hamsters, and NHPs. We provided evidence that RQ3013 is a promising vaccine candidate against SARS-CoV-2 VOCs, adding support for its clinical development.

## Results

### Antigen design and characterization

RQ3013 encodes a near full-length S protein lacking the eighteen cysteine-rich residues at the extreme C-terminus. The native signal peptide and the transmembrane domain are kept in the immunogen. Compared to ancestral S protein from the wild-type SARS-CoV-2, RQ3013 immunogen has all the mutations derived from the B.1.1.7 variant with additional K417N, E484K, and A701V mutations found in the B.1.351 variant. Seven mutations also exist in the Omicron variant B.1.1.529. Unlike the S2P design used in BNT162b2 and mRNA-1273^5, 6^, the RQ3013 immunogen does not have those two proline substitutions. Instead, the furin cleavage site ^682^RRAR^685^ is mutated to GSAS to prevent protease-mediated proteolysis (Fig. 1a). Robust expression of S protein was confirmed in HEK293A cells upon transfection of RQ3013 mRNA formulated in LNP or mixed with a transfection reagent. Western blot with the cell lysate under denaturing conditions detected a single band in the cell lysate corresponding to the uncleaved S protein with an apparent molecular mass of 170kDa (Fig. S1).

**Fig. 1.**
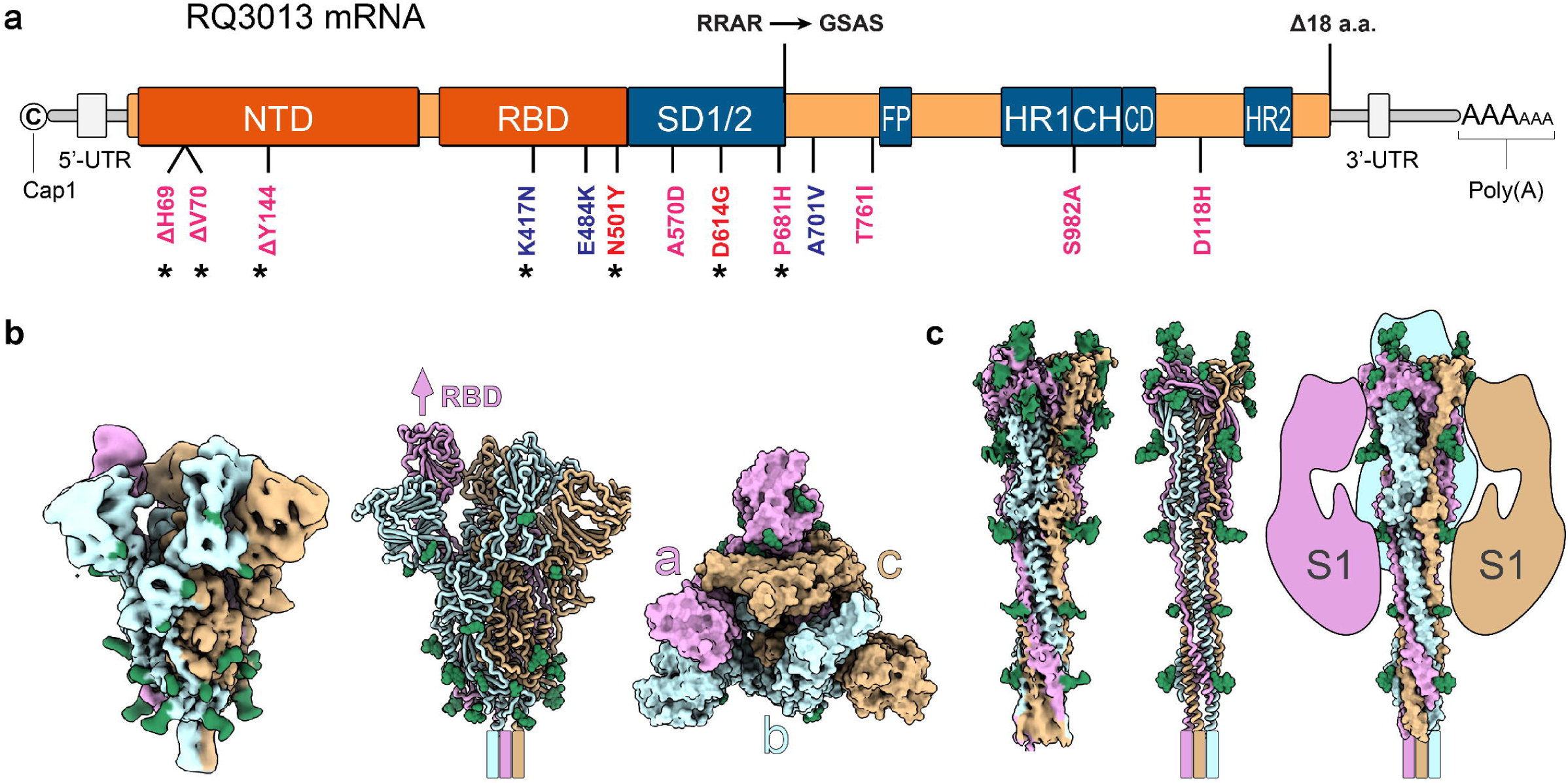
Vaccine design of RQ3013 and structural characterization of the expressed antigens. **a** An illustration of RQ3013 mRNA. Mutations compared to the wild-type S were indicated. Pink ones were derived from the SARS-CoV-2 B.1.1.7 variant, Blue from B.1.351, and Red shared by both variants. Starred also existed in B.1.1.529. The furin cleavage site RRAR was replaced by GSAS.The last 18 amino acids were deleted. Abbreviations: UTR, untranslated region; NTD, N-terminal domain; RBD, receptor-binding domain; SD1/2, subdomain 1 and 2; FP, fusion peptide; HR1, heptad repeat 1; CH, central helix; CD, connector domain; HR2, heptad repeat 2. **b** The cryo-EM map and the refined atomic model of the S antigen in the prefusion conformation. One RBD in the upright state was indicated. **c** The cryo-EM map and the refined model of the postfusion S antigen. The unseen S1 domains were sketched. Protomers were colored in pink, cyan, and orange, and glycosylations were colored in green.

To characterize the S protein encoded by RQ3013, we overexpressed and affinity-purified the antigen and investigated its 3D structure by single-particle cryo-EM analysis. Initial 2D classification revealed two distinct populations corresponding to prefusion and postfusion S, from which two mass density maps were produced at a nominal resolution of 3.87 Å and 3.49 Å, respectively (Fig. S2). The refined models revealed homotrimeric S in the prefusion or postfusion state. The prefusion S has one RBD in the upright open conformation. Further 3D classification in the population did not reveal other conformations (Fig. 1b). This dominant one-RBD up form resembles S from the Omicron variant^7^ and differs from the ancestral S, which takes a mixture of open (RBD up) and closed (RBD down) conformations.

A portion of purified S proteins also existed in a postfusion state (Fig. 1c and Fig. S2). No cleavage was observed in the sample used for the cryo-EM analysis. This indicates that the structural transition of SARS-CoV-2 S protein from pre-to postfusion may occur in the absence of cleavage, which has been reported for other class I fusion proteins^8^. Whether such transition occurs in vivo, or to what extent, remains to be determined as the postfusion structure could result from the purification process in the presence of detergent.

## Immunogenicity of RQ3013 in BALB/c mice

To assess the immunogenicity of RQ3013 in BALB/c mice. The two groups of mice (n = 18) were immunized intramuscularly with 2 µg or 5 µg RQ3013 on day 0 (Fig. 2a). A control group of mice (n = 18) received PBS as the placebo. All groups were boosted on day 21. No local inflammation or other adverse effects were observed throughout the experiment. We evaluated sera collected on day 28 for binding IgG to RBD and S. Data from enzyme-linked immunosorbent assay (ELISA) showed that the 2 µg RQ3013 induced in mice equally high titers of IgG binding to Wild-type RBD and S from the B.1.1.7, B.1.351, and B.1.617.2 variants (Fig. 2b). There was no difference between the low-dose group (2 μg) and the high-dose group (5 μg)(Fig. 2b). Next, we performed the lentiviral pseudovirus neutralization assay to evaluate the levels of neutralizing antibodies (NAb) induced by RQ3013. Sera on day 28 exhibited a balanced cross-activity against Wild-type, B.1.1.7, B.1.351, and B.1.617.2 virus (Fig. 2c). In mice sera from the low-dose group, there existed similar levels of NAbs against Wild-type (geometric mean titer(GMT) 469), B.1.1.7 (GMT 400), and B.1.351 (GMT 664). For B.1.617.2, there was a 2 to 3-fold reduction in NAb titer (GMT 218). In the 5-µg dose group, titers increased for all tested variants except B.1.351 (Fig. 2c). To investigate whether RQ3013 activates a T cell response in mice, we collected peripheral blood mononuclear cells (PBMCs) on day 41 and re-stimulated them with the S peptide mix. Using an enzyme-linked immunosorbent spot (ELISPOT) assay, high levels of IFN-γ secreting in Th1 cells were detected in all RQ3013-vaccinated mice but not in the mice received PBS (Fig. 2d).

**Fig. 2.**
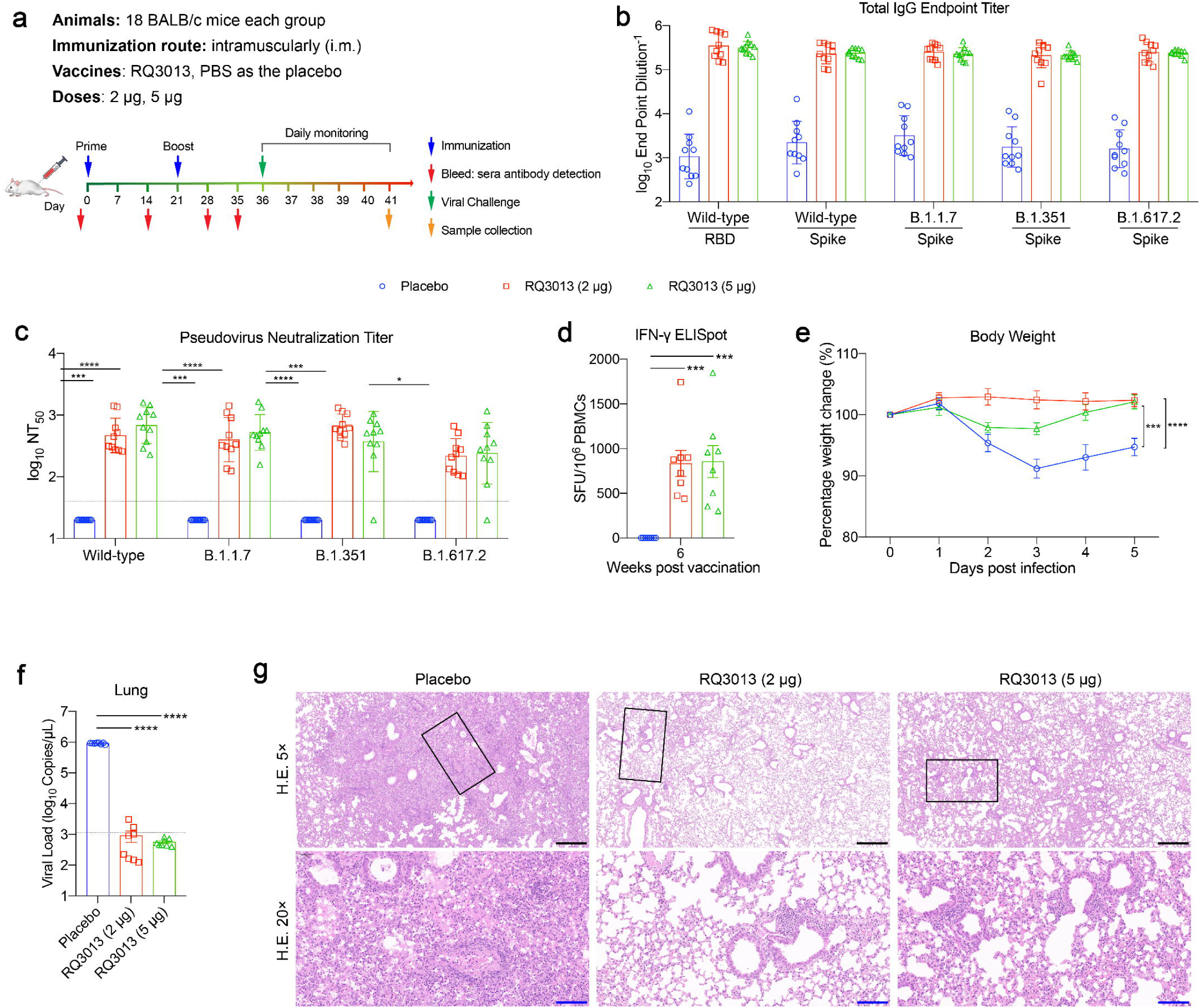
Immunogenicity and protective efficacy of RQ3013 in BALB/c mice. **a** The immunization scheme of RQ3013 in mice. Mice (n = 18 per group) were intramuscularly immunized with PBS(Placebo, blue circle) or low dose (2 μg, red square) or high dose (5 μg, green triangle) of RQ3013. Time points of vaccination, bleeding, viral challenge, and sera collection are indicated by arrows. **b** ELISA analysis of binding IgG to the wild-type RBD antigen and S proteins from SARS-CoV-2 variants with sera collected on day 28. Values are GMT mean ± SD. **c** Neutralizing antibody titers in sera collected on day 28, analyzed by the lentiviral luciferase-based pseudovirus assay. The black dashed line indicates the assay’s detection limit (reciprocal titer of 40). Any measurement below the detection limit was assigned a value of half the detection limit for plotting and statistical purposes. Values are GMT mean ± SD. **d** Summed IFN-γ ELISpot responses in PBMCs restimulated with peptides spanning the entire S. The PBMCs were collected on day 41. SFU, spot-forming units. Data are presented as mean ± SEM. **e** Body weight changes of BALB/c mice following infection with the variant B.1.351 (n = 6). Data are presented as mean ± SEM. **f** Viral RNA loads in mice lung tissues at 5 dpi, determined by qRT-PCR. The black dashed line indicates the assay’s detection limit (1146 Copies/µL). Data are presented as mean ± SEM. **g** Histopathological examinations in mice lungs at 5 dpi. Lung tissue was collected and stained with hematoxylin and eosin. Black scale bar, 400 μm; blue scale bar, 100 μm. Statistical analyses were carried out by Student’s t-test when two groups were analyzed, and by ANOVA when more than two groups were analyzed (*P < 0.05; **P < 0.005; ***P < 0.001; ****P < 0.0001).

### Efficacy of RQ3013 in BALB/C mice

Wild-type BALB/c mice, which do not exhibit infection by the wild-type SARS-CoV-2 virus, are susceptible to variants B.1.1.7 and B.1.351^9^. We, therefore, evaluated the protection efficacy of RQ3013 in BALB/c mice. The immunized mice were challenged intranasally with the variant B.1.351 (1×10^5^ TCID_50_) 15 days after the boost. The lung tissues were collected at 5 days post-infection (dpi) for viral load determination (by qRT-PCR) and histopathological analysis (Fig. 2a). Compared to the PBS control, both 2-μg and 5-μg doses of RQ3013 prevented weight loss of mice beginning at 2 dpi (Fig. 2e). All mice in the placebo group developed a high load of viral sgRNAs (∼10^6^ sgRNA copies/µL) in the lungs, which were not detectable in both groups of immunized mice (Fig. 2f). Accordingly, lung pathology showed intact alveolar structures in immunized mice, while mice in the control group developed pathological characteristics of typical lung lesions (Fig. 2g). These results demonstrated that RQ3013 could effectively prevent replication of the B.1.351 variant in the lower respiratory tract and provides rapid protection in BALB/c mice from lung lesions.

### Immunogenicity and protection of RQ3013 in K18-hACE2 transgenic mice

In the K18-hACE2 transgenic mouse model, B.1.1.7 and B.1.351 are 100 times more lethal than the original virus^10^. We next assessed the immunogenicity and efficacy of RQ3013 in K18-hACE2 transgenic mice. Both 2-μg and 5-μg doses of RQ3013 elicited high titers of RBD-and S-specific IgG antibodies in sera of K18-hACE2 mice at 7 days post-boost in a dose-dependent manner (Fig. 3a, b). We also measured inhibition of cell entry of viruses pseudotyped with the wild-type and B.1.1.529 S protein. 5-μg dose elicited neutralizing antibodies in all mice for Wild-type (GMT 434) and B.1.1.529 (GMT 173) (Fig. 3c). We next challenged the mice with the variant B.1.351 on day 61 (1×10^3^ PFU). Mice in the control group showed weight losses at 4 dpi, which were prevented in both dose groups of RQ3013-vaccinated mice (Fig. 3d). Mice were euthanized at 4 dpi, and a large amount of viral RNAs was detected in the lung (10^6^ copies per μg of RNA), trachea (10^5^ copies per μg of RNA), and brain (10^9^ copies per μg of RNA) of the control mice. RQ3013 effectively prevented viral replication in these tissues. A decrease in viral load (> 3 logs) in the lung was observed in all RQ3013-vaccinated mice. Viral replication was completely prevented in the brain except for two mice in the 2-μg dose group (Fig. 3e).

**Fig. 3.**
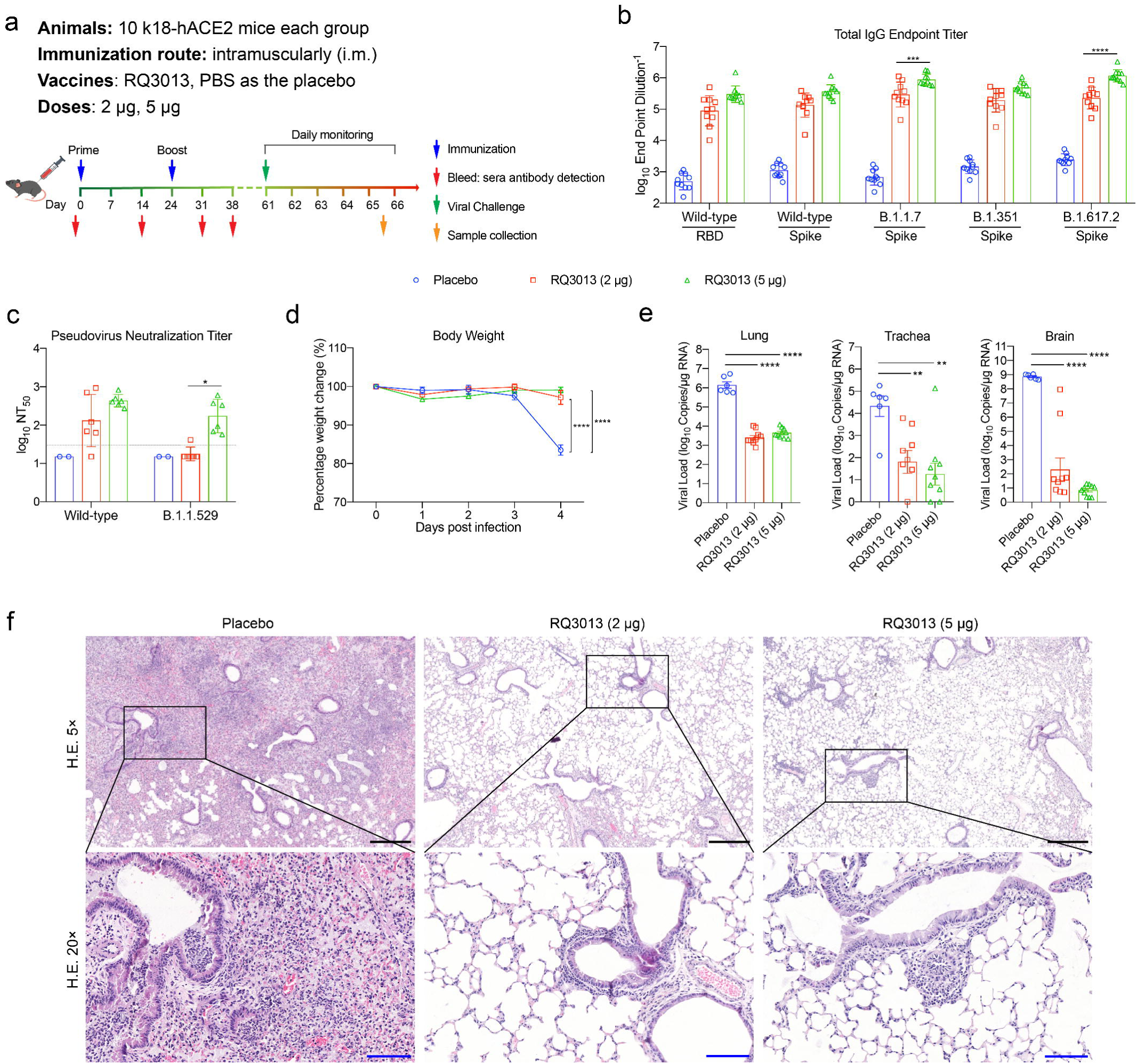
Immunogenicity and protective efficacy of RQ3013 in K18-hACE2 transgenic mice. **a** The scheme of mice immunization. Mice (n = 10 per group) were intramuscularly immunized with PBS (Placebo, blue circle) or low dose (2 μg, red square) or high dose (5 μg, green triangle) of RQ3013. Time points of vaccination, bleeding, viral challenge, and sample collection are indicated by arrows. **b** ELISA analysis of binding IgG to the wild-type RBD antigen and S proteins from SARS-CoV-2 variants with sera collected on day 31. Values are GMT mean ± SD. **c** Neutralizing antibody titers in day-31 sera, analyzed by the lentiviral luciferase-based pseudovirus assay. Values are GMT mean ± SD. **d** Body weight changes of K18-hACE2 mice following infection with the variant B.1.351. Data are presented as mean ± SEM. **e** Viral RNA loads in the mice lung, trachea, and brain tissues at 4 dpi, determined by qRT-PCR. Data are presented as mean ± SEM. **f** Histopathological examinations in mice lungs at 4 dpi. Lung tissues were collected and stained with hematoxylin and eosin. Black scale bar, 400 μm; blue scale bar, 100 μm. Statistical analyses were carried out by Student’s t-test when two groups were analyzed, and by ANOVA when more than two groups were analyzed (ns, Not Significant; *P < 0.05; **P < 0.005; ***P < 0.001; ****P < 0.0001).

Lung histology revealed severe lung lesions in PBS-immunized mice characterized by the diffuse thickening of the alveolar septae, mononuclear cell infiltration, and vascular congestion. In contrast, mice vaccinated with RQ3013 did not develop pathological changes, showing good protection against the B.1.351 variant of RQ3013 in K18-hACE2 mice (Fig. 3f)

### Immunogenicity of RQ3013 in hamsters

The Syrian hamster having the ACE2 receptor is highly susceptible to SARS-CoV-2 and develops similar pneumonia to that in COVID-19 patients, making them a suitable model for evaluating vaccines^11^. Three groups of hamsters were vaccinated on day 0 and day 21 with PBS or RQ3013 (5 or 25 µg) (Fig. 4a), and sera were collected and evaluated for immunogenicity on day 28. High titers of binding antibodies against RBD and S (wild-type and B.1.617.2) were detected (Fig. 4b). We monitored the level of S-specific IgG over the course of 33 weeks post-immunization, and a reduction in titer was observed in the first 13 weeks and then sustained at a high level for the rest time (Fig. S3). The pseudovirus neutralization assay showed that high titers of NAbs against Wild-type, B.1.1.7, B.1.351, B.1.617.2, and B.1.1.529 were elicited in vaccinated mice in a dose-dependent manner, and the ability to neutralize the B.1.1.529 variant is the lowest (Fig. 4c). The NAb titer was also measured with the CPE-based colorimetric live virus micro-neutralization assay. Compare to B.1.617.2, there was a four-fold drop in the level of neutralizing antibodies against B.1.1.529. Nonetheless, the titers were still significant with GMT 160 and 242 for 5 μg and 25 μg doses, respectively (Fig. 4d). We also collected PBMCs and re-stimulated them with the S1 and S2 peptide pools. Using an enzyme-linked immunosorbent spot (ELISPOT) assay, we detected high levels of IFN-γ secreting cells in RQ3013-immunized hamsters in a dose-dependent manner (Fig. 4e). These data confirmed that the mRNA vaccine could induce a strong T cell response in hamsters.

**Fig. 4.**
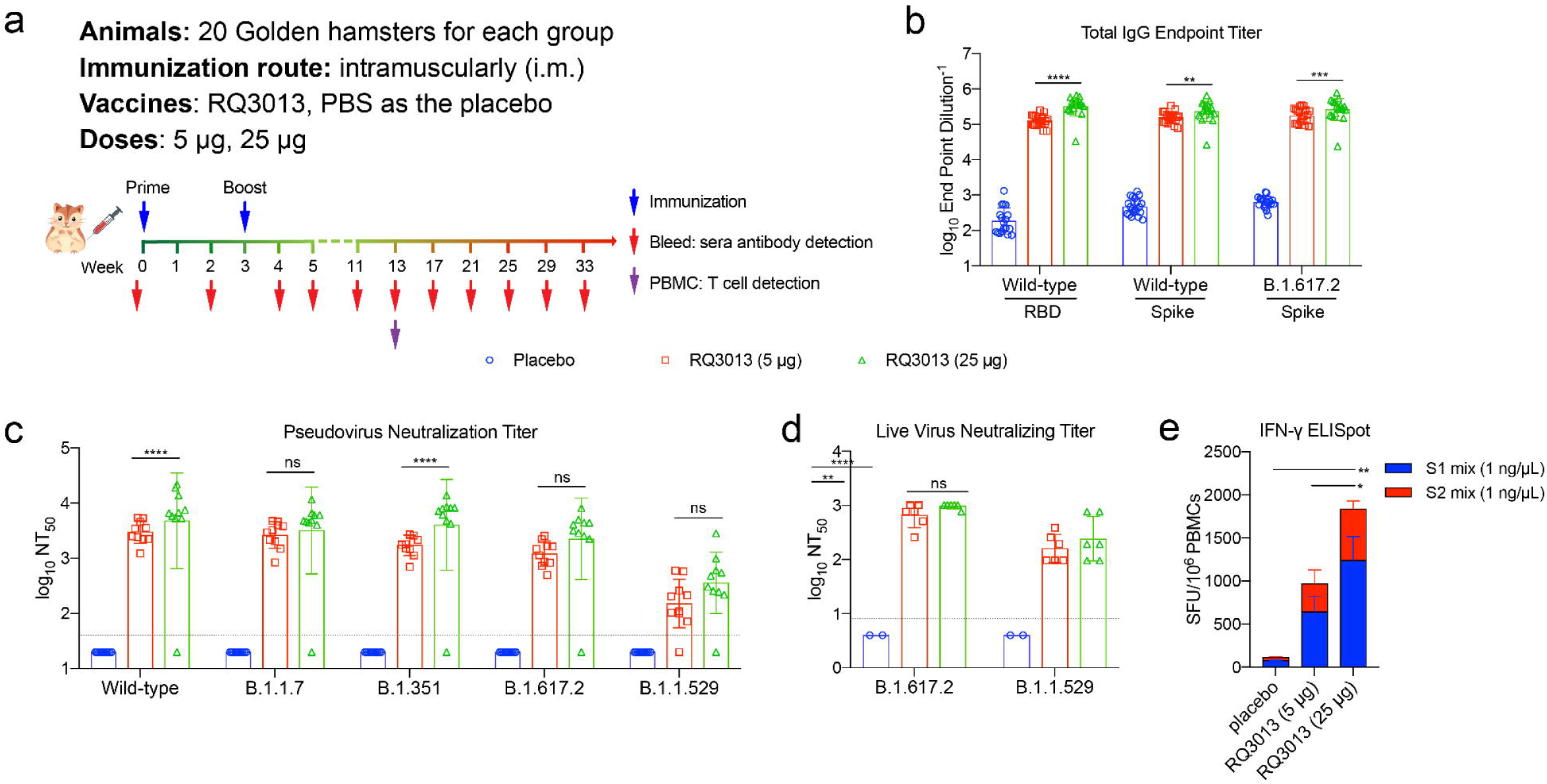
Immunogenicity of RQ3013 in hamsters. **a** The immunization scheme of hamsters by RQ3013. Hamsters (n = 20 per group) were intramuscularly immunized with PBS (Placebo, blue circle) or low dose (5 μg, red square) or high dose (25 μg, green triangle) of RQ3013. Time points of vaccination, bleeding, viral challenge, and PBMCs collection are indicated by arrows. **b** Binding IgG to the wild-type RBD and S proteins from the wild-type and B.1.617.2 viruses, analyzed with sera collected in week 4 by ELISA. Values are GMT mean ± SD. **c** Neutralizing antibody titers in week-4 sera, analyzed by the lentiviral luciferase-based pseudovirus assay. The black dashed line indicates the assay’s limit of detection (reciprocal titer of 40). Any measurement below the detection limit was assigned a value of half the limit of detection for plotting and statistical purposes. Values are GMT mean ± SD. **d** The CPE-based live virus micro-neutralization assay of week-5 sera against SARS-CoV-2 B.1.617.2 and B.1.1.529. The black dashed line indicates the assay’s detection limit (reciprocal titer of 8). Any measurement below the detection limit was assigned a value of half the limit of detection for plotting and statistical purposes. Values are GMT mean ± SD. **e** The IFN-γ ELISpot assay with PBMCs obtained in week 13 and restimulated with overlapping peptide pools of S1 and S2 domains. SFU, spot-forming units. Data are presented as mean ± SEM. Statistical analyses were carried out by Student’s t-test when two groups were analyzed, and by ANOVA when more than two groups were analyzed (ns, Not Significant; *P < 0.05; **P < 0.005; ***P < 0.001; ****P < 0.0001).

### Immunogenicity and protection of RQ3013 in rhesus macaques

We next evaluated the immunogenicity of RQ3013 in rhesus macaques, an NHP model susceptible to SARS-CoV-2 infection^12^. Two groups of rhesus macaques (n = 4/group, 2 of each sex, ages 6 to 8 years) were primed with 30 µg or 100 µg of RQ3013 via intramuscular administration and boosted on day 21 (Fig. 5a). A third group (n = 4) received PBS as the control (Fig. 5a). All macaques receiving RQ3013 developed RBD- and S variants-specific IgG in sera collected on days 28 or 35 (Fig. 5b). Peudovirus assay displayed a broad-spectrum ability to neutralize SARS-CoV-2 variants, including B.1.1.529, consistent with data from the mouse and hamster model (Fig. 5c). Using sera on day 42, we performed the plaque reduction neutralization test (PRNT) with four live viruses. In the 30-µg dose group, the PRNT_50_ GMT of neutralizing antibodies was 1081, 1775, 304, and 867 for Wild-type, B.1.351, B.1.617.2, and B.1.1.529, respectively. There were no significant changes in the 100-µg dose group, except that the GMT increased to 3198 for the B.1.351 variant (Fig. 5d). The PRNT_50_ data agree reasonably well with the antigen design of RQ3013 which is more specific to B.1.351 than Wild-type and B.1.617.2. Interestingly, the level of cross-neutralizing antibody against B.1.1.529 is encouragingly high in NHP, near the level for the Wild-type virus (Fig. 5d).

**Fig. 5.**
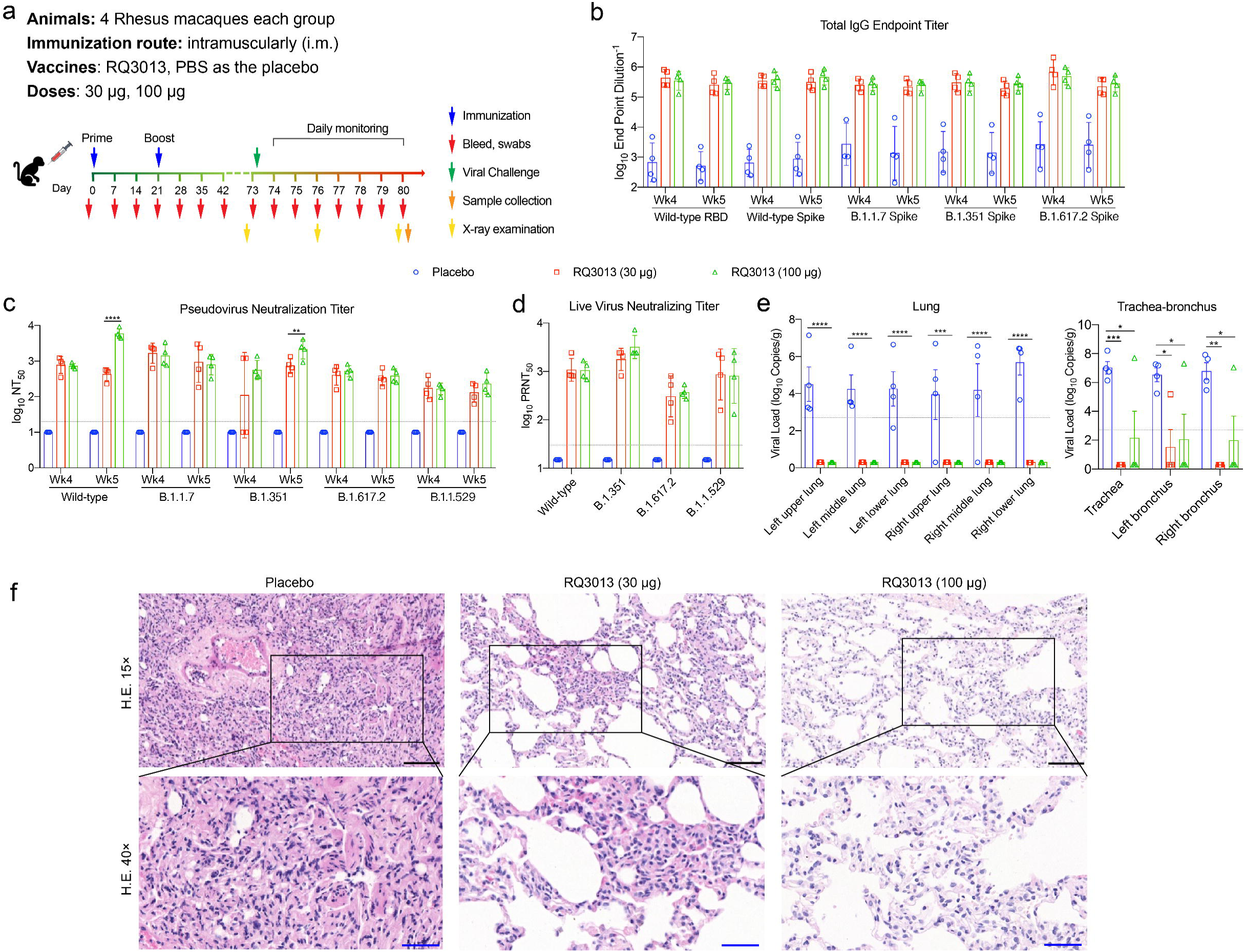
Immunogenicity and protection of RQ3013 in NHP rhesus macaques. **a** The immunization scheme of RQ3013. Macaques (n = 4 per group) were vaccinated on days 0 and 21 through the intramuscular route with PBS (Placebo, blue circle) or low dose (30 μg, red square) or high dose (100 μg, green triangle) of RQ3013. Time points of vaccination, bleeding, viral challenge, sample collection, and X-ray examination are indicated by arrows. **b** Binding IgG to the wild-type RBD antigen and S proteins from variants, analyzed with sera collected on days 28 (Wk4) and 35 (Wk5) by ELISA. Values are GMT mean ± SD. **c** Neutralizing antibody titers in sera from days 28 and 35, analyzed by the lentiviral luciferase-based pseudovirus assay. The black dashed line indicates the assay’s detection limit (reciprocal titer of 20). Any measurement below the detection limit was assigned a value of half the limit of detection for plotting and statistical purposes. Values are GMT mean ± SD. **d** The PRNT-based neutralization assay of day-42 sera against SARS-CoV-2 variants. The black dashed line indicates the assay’s limit of detection (reciprocal titer of 30). Any measurement below the detection limit was assigned a value of half the detection limit for plotting and statistical purposes. Values are GMT mean ± SD. **e** Viral RNA loads in aminal lungs and trachea-bronchus at 7 dpi, determined by qRT-PCR. The black dashed line indicates the assay’s detection limit (500 Copies/g). Any measurement below the detection limit was recorded as ‘2’ for plotting and statistical purposes. Data are presented as mean ± SEM. **f** Histopathological examinations of aminal lungs at day 7 dpi. Lung tissues were collected and stained with hematoxylin and eosin. Black scale bar, 100 μm; blue scale bar, 60 μm. Data are presented as mean ± SEM. Statistical analyses were carried out by Student’s t-test when two groups were analyzed, and by ANOVA when more than two groups were analyzed (*P < 0.05; **P < 0.005; ***P < 0.001; ****P < 0.0001).

At 7 weeks following the second vaccination, rhesus macaques were challenged with the Wild-type virus (1×10^5^ TCID_50_). All animals exhibited no noticeable behavioral differences and no apparent changes in body weight and body temperature during the experimental period (Fig. S4a, b).

In the low-dose group of animals, viral RNA was not detected since day 1 post-infection in the nasal and anal swabs and became absent at 6 dpi in the oropharyngeal swabs. A similar trend was observed for the high-dose group. In contrast, the PBS-vaccinated animals experienced a continuous viral replication as seen in anal and oropharyngeal swabs. Viruses were not detected in blood samples of all animals (Fig. S4c-f). These data showed rapid control of viral replication within 2 days in both the upper and lower airways conferred by RQ3013 vaccination.

After sacrifice, no viral RNA was detected in the lungs of all RQ3013-vaccinated animals. One from each dose group has viral RNA in trachea-bronchus detected. A high load of viruses was seen in the lung, trachea, and bronchus of animals in the control group (Fig. 5e).

X-ray images showed no noticeable lung shadow in all infected animals (Fig. S5).

Pathological changes characterized by viral pneumonia and pulmonary fibrosis were moderate in the low-dose group and generally mild in the high-dose group, significantly mitigated compared with the control group (Fig. 5f).

### Safety evaluation of RQ3013 in nonhuman primates

We evaluated the safety of RQ3013 in NHP cynomolgus macaques. Fifty cynomolgus macaques were divided into five groups (n = 10), each on day 0 received intramuscularly one dose of PBS or RQ3013 (60 µg and 240 µg) or empty LNPs (1.2 mg or 4.8 mg of total lipid) (Fig. 6a). All groups were double-boosted on days 14 and 28. Neither fever nor weight loss was observed in all animals (Fig. S6). The blood samples of all animals were collected at different time points for hematological and biochemical analysis. All animals displayed no appreciable changes in hematological indices and the percentage values of lymphocyte subsets, including CD3^+^, CD4^+^, and CD8^+^ T cells (Fig. S7 and Fig. 6b). For key cytokines, no notable changes in Th1 cytokines (interferon-γ, tumor necrosis factor α, interleukin-2) or Th2 cytokines (interleukin-4, 5) secretion was observed in all animals (Fig. 6c and Fig. S8). The IL-6 level increased following each injection of either RQ3013 or empty LNP in a dose-dependent manner. All recovered in a week. (Fig. S8). We detected high levels of IFN-γ secreting Th1 cells and interleukin-4 (IL-4) secreting Th2 cells in animals immunized with RQ3013 but not PBS or empty LNP (Fig. S9).

**Fig. 6.**
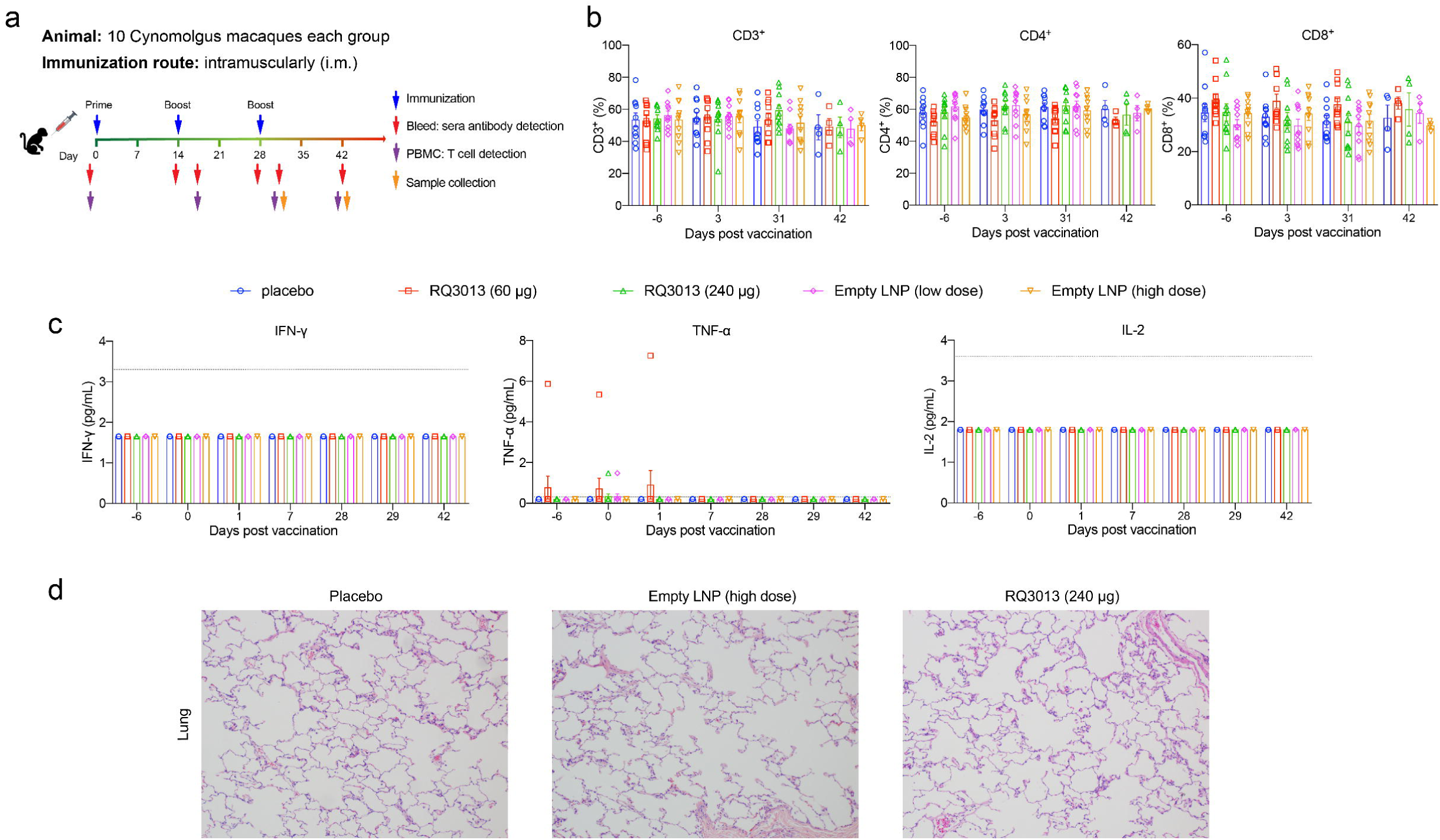
Safety evaluation of RQ3013 in NHP cynomolgus macaques. **a** The immunization scheme of macaques. Macaques (n = 10 per group) were immunized three times on days 0, 14, and 28 through the intramuscular route with one of the following substances: PBS (Placebo, blue circle), low dose RQ3013 (60 μg, red square), high dose RQ3013 (240 μg, green triangle), low dose (purple diamond) and high dose (orange inverted triangle) of empty LNP. Time points of vaccination, bleeding, and sample collection are indicated by arrows. (**b**, **c**) Hematological analysis in all five groups of macaques. Lymphocyte subset percents, including CD3^+^, CD4^+,^ and CD8^+^, and cytokines IFN-γ, TNF-α, and IL-2 were monitored on days indicated. Values are mean ± SEM. The black dashed line indicates the assay’s detection limit (IFN-γ, 3.3 pg/mL; TNF-α 0.3 pg/mL; IL-2, 3.6 pg/mL). Any measurement below the detection limit was assigned a value of half the limit of detection for plotting and statistical purposes. Values are mean ± SEM. **d** Histopathological evaluations in lungs from three groups of macaques on day 31. Lung tissues were collected and stained with hematoxylin and eosin. Statistical analyses were carried out by ANOVA when more than two groups were analyzed.

Finally, three groups of animals (PBS, high-dose groups for RQ3013 and empty LNP) were euthanized on day 31. Lungs, brains, hearts, livers, spleens, and kidneys were harvested for histopathological analysis and safety assessment of RQ3013. Vaccine-associated immunopathologic changes were not observed in any of the sections examined in these tissues of all animals (Fig. 6d and Fig. S10). Together, these data provided evidence that a high dose application of RQ3013 is safe in NHP, supporting the development of clinical trials.

## Discussion

The mRNA vaccines mRNA-1273 and BNT162b2 were developed at the beginning of the COVID-19 pandemic and have shown greater than 90% efficacy during the early phase of the pandemic^13^. However, the protection against new infections has dropped considerably for both vaccines with the continuous emergence of SARS-CoV-2 variants carrying mutations in the spike protein, the antigenic target of the mRNA vaccines. Under this background, we developed an mRNA vaccine candidate, RQ3013, to provide broad-spectrum protection against VOCs. The immunogen encoded in RQ3013 is the S protein with the last 18 residues truncated. It carries all the mutations found in the B.1.1.7 variant. In addition, it has K417N, E484K, and A701V associated with the B.1.351 variant. B.1.1.529 is phylogenetically closer to B.1.1.7, and they share the ΔH69/V70, ΔY144, D614G, N501Y, and P681H mutations across the S protein. B.1.1.529 spike also harbors a K417N mutation in RBD, which, together with N501Y, has been implicated in causing immune evasion^14^. Our results in various animal models showed RQ3013 elited uniformly high levels of NAb against Wild-type, B.1.1.7, and B.1.351. In general, it’s more effective against B.1.351 than against Wild-type, agreeing with its antigen design. Conversely, an 8 to 10-fold reduction in neutralizing activity against B.1.351 compared to Wild-type was observed in BNT162b2-elicited human sera^15, 16^. RQ3013 is less competent in eliciting neutralizing activities against B.1.617.2 and B.1.1.529, which are generally 3 to 5-fold lower than against Wild-type in mice and hamsters. Notably, In NHP rhesus macaques vaccinated with two doses of RQ3013, the NAb titer (PRNT_50_) against B.1.1.529 in sera is almost to the same level as against Wild-type. This is in contrast with vaccines currently in use, where a drastic reduction in neutralizing activity against Omicron was observed^17^. Our data demonstrated that RQ3013 could elicit relatively balanced cross-neutralizing antibody responses.

In addition to variant-related mutations, we also changed the furin cleavage site in S to eliminate post-translational proteolysis, thus stabilizing S in the prefusion state. However, structural characterization revealed trimeric S in both prefusion and postfusion states. Although we could not locate the S1 subunit in the postfusion structure due to its flexibility, it most likely remained in the structure. A previous study used the same strategy to determine the S structure only to report the prefusion conformation^6^. The structural transition observed in our study may be induced by harsh detergent conditions during purification. It is evident that prevention of proteolysis can stabilize the prefusion conformation but not to the extent conferred by proline substitutions^18^. On the other hand, the exposure of the S2 subunit may be beneficial for vaccine development against SARS-CoV-2. Since S2 is more conserved than S1, antibodies targeting S2 have shown great cross-neutralization activity against β-coronaviruses^19, 20^. Therefore, it is possible that the S2-binding antibodies may partly contribute to the broad spectrum of RQ3013.

The protection efficacy of RQ3013 was demonstrated in BALB/c and K18-hACE2 mice and NHP rhesus macaques. Two doses of RQ3013 effectively protected mice from infection by B.1.351. Although RQ3013 elicited a lower titer of NAb against the wild-type SARS-CoV-2, two doses of this vaccine protected the upper and lower respiratory tract of NHPs after the challenge with the wild-type virus.

In summary, we have conducted a broad-spectrum assessment of RQ3013 immunogenicity, efficacy, and safety, providing support for the evaluation of this vaccine in clinical trials.

## Materials and Methods

### Facility and ethics statement

All experiments with live SARS-CoV-2 viruses were performed in the enhanced biosafety level 3 (P3+) facilities in Guangzhou Institute of Respiratory Health (GIRH), Wuhan Institute of Virology (WIV), and the Institute of Medical Biology, Chinese Academy of Medical Sciences. All experiments with mice, hamsters, and macaques were carried out in accordance with the Regulations in the Guide for the Care and Use of Laboratory Animals of the Ministry of Science and Technology of the People’s Republic of China.

The female BALB/c mice (6- to 8-week old) and female K18-hACE2 Transgenic Mice (6- to 8-week old) experiments with SARS-CoV-2 challenge were conducted under animal biosafety level 3 (ABSL3) facilities in Guangzhou Institute of Respiratory Health (GIRH) and Wuhan Institute of Virology (WIV) approved by the National Health Commission of the People’s Republic of China respectively.

The 6- to 8-year-old male or female rhesus macaque experiments were performed in the animal biosafety level 4 (ABSL-4) facility at Wuhan Institute of Virology (WIV), Hubei, China. All animal procedures were approved by the Institutional Animal Care and Use Committee of the Institute of Medical Biology, Chinese Academy of Medical Science.

### Cell lines and antibodies

HEK 293F cells were grown in FreeStyle Media (Gibco-Thermo Fisher Scientific) and transiently transfected using polyethylenimine (PEI) (Polysciences, Inc.) in an 8% CO_2_ environment at 37°C. HEK 293A and Vero E6 cells were maintained in high glucose DMEM(GIBCO) supplemented with 10% FBS(GIBCO) and 1% penicillin/streptomycin (P/S) (GIBCO) in a 5% CO_2_ environment at 37°C.

The following primary antibodies were used in this study: anti-SARS-CoV-2 (2019-nCoV) Spike Antibody (40589-T62, Sino Biological), and anti-GAPDH Antibody (60004, Proteintech). The secondary antibodies used were Peroxidase AffiniPure Goat Anti-Rabbit IgG (H+L) (111-035-003, Jackson ImmunoResearch), Peroxidase AffiniPure Goat Anti-Mouse IgG (H+L) (115- 035-003, Jackson ImmunoResearch), goat anti-Syrian hamster IgG HRP (abcam Cat: ab6892) and goat anti-monkey IgG HRP (Invitrogen Cat: PA1- 84631).

### Virus titration

Virus titrations were performed by endpoint titration in Vero E6 cells. Cells were inoculated with 10-fold serial dilutions of cell supernatant. One hour after inoculation of cells, the inoculum was removed and replaced with 100 μL of DMEM supplemented with 2% FBS, 1 mM L-glutamine, 100 IU/mL penicillin, and 100 μg/mL streptomycin. Three days after inoculation, CPE was scored, and the TCID50 was calculated.

### Preparation of modified mRNAs and LNP

Production of RQ3013 mRNA and LNP used in this study was conducted under current good manufacturing practice (cGMP). The optimized DNA template for RQ3013 RNA was cloned into the plasmid with backbone elements (T7 promoter, 5′ and 3′ UTR, poly(A) tail). RQ3013 mRNA was in vitro-transcribed by T7 RNA polymerase in the presence of the Cap1 analog (B8176, APExBIO) and nucleotides with a global substitution of uridine with pseudouridine (Ψ, APExBIO). RNA was purified through two chromatographic procedures. RNA integrity was assessed by microfluidic capillary electrophoresis, and the concentration, pH, osmolality, and endotoxin level were determined. Purified RNA was formulated into lipid nanoparticles containing four lipids (an ionizable lipid, PEG2000-DMG, DSPC, and cholesterol) as previously described. Particle sizes were measured using dynamic light scattering on a Malvern Zetasizer Nano-ZSP (Malvern).

### Protein expression of mRNAs and LNPs

The protein expression of mRNAs was tested in HEK 293A cells. mRNA transfection was carried out using lipofectamine 2000 (lipo2K) at a ratio of 1:2 (mRNA: lipo2K). For LNP, 5 μg of LNPs were incubated with HEK 293A cells in one well of a 12-well plate for 18 hours, and cells and media were collected for protein analysis. Protein expression was detected by western blotting, similar to previously. Briefly, cells were collected and rinsed with PBS and lysed by lysis buffer (20mM Tris-HCl (pH = 7.4), 150mM NaCl, 3mM MgCl_2_, 1% Triton X-100) supplemented with protease inhibitor. All samples were mixed with SDS loading buffer, separated in a 4-20% gradient SDS-Gel, and transferred to PVDF membranes (ThermoFisher) by Trans-Blot Turbo Transfer System (BioRad). The blots were blocked with 5% non-fat dry milk in PBST and then incubated with appropriate primary antibodies. Signals were detected with HRP-conjugated secondary antibodies and an enhanced chemiluminescence (ECL) detection system.

### Cloning, expression, and preparation of the RQ3013 encoded Spike proteins

The gene encoding the RQ3013 was fused with a C-terminal twin Strep-tag (LEVLFQGPSGS WSHPQFEK GGGSGGGSGGSA WSHPQFEK) and cloned into a mammalian cell expression vector pcDNA3.1. The resulting plasmid, pcDNA3.1-RQ3013-Twinstrep, was transformed into HEK 293F cells using polyethylenimine (PEI) in FreeStyle Media (Gibco-Thermo Fisher Scientific).

To purify the full-length S protein, the transfected cells were harvested at a density of ∼4-5×10^6^/mL by centrifugation (1,000 × g at 4°C for 30 min). The cell pellet was washed with cold PBS, resuspended in a cold lysis buffer containing Buffer A (100 mM Tris-HCl, pH 8.0, 150 mM NaCl, 1 mM EDTA) and 1% NP- 40 (w/v), 1 × EDTA-free complete protease inhibitor cocktail (Roche, Basel, Switzerland), and incubated at 4°C for one hour.

The sample preparation steps are described elsewhere (Science. 2020 Jul 21: eabd4251). Briefly, after clarifying spin at 30,000 × g for 60 min at 4°C, the supernatant was loaded on a strep-tactin (IBA Lifesciences, Germany) column equilibrated with the lysis buffer. The S protein was eluted by Buffer A containing 0.02% NP-40 and 50 mM biotin and further purified by gel filtration chromatography on a Superose 6 10/300 column in a buffer containing 25 mM Tris-HCl, pH 7.5, 150 mM NaCl, and 0.02% NP-40.

### Cryo-EM grids preparation, data collection, and image processing

To prepare cryo-EM grids, 3.5 μL of freshly purified spike proteins at ∼0.3 mg/mL was applied to Quantifoil R1.2/1.3 Au holey carbon grids supported by a continuous thin layer of carbon film (Quantifoil Micro Tools GmbH, Germany), which had been glow discharged with a PELCO easiGlow^TM^ Glow Discharge Cleaning system (Ted Pella, Inc., Redding, CA) for 30 s at 15 mA. Grids were immediately plunge-frozen in liquid ethane using a Vitrobot Mark IV (Thermo Fisher Scientific). Excess protein was blotted away using grade 595 filter paper (Ted Pella, Inc.) with a blotting time of 4 seconds, and a blotting force of -12 at 4°C in 100% humidity.

The cryo-EM grids were loaded onto a Thermo Fisher Scientific Titan Krios G3i transmission electron microscope equipped with a Gatan GIF Quantum energy filter (slit width 20eV) and operated at 300 kV for data collection. All the cryo-EM images were automatically recorded by a post-GIF Gatan K3 direct electron detector in the super-resolution counting mode using EPU software (version 2.8.2.10REL) with a nominal magnification of 81,000 × in the EFTEM mode, which yielded a super-resolution pixel size of 0.55 Å on the image plane, and with a defocus ranging from -1.5 to -2.5 μm. Each micrograph stack was dose-fractionated into 40 frames with a total electron dose of ∼ 50 e^-^/ Å^2^ and a total exposure time of 3 s. For the cryo-EM dataset, 1,474 micrographs from a total of 4,001 micrographs were selected for further processing.

For image processing, drift and beam-induced motion correction were applied on the super-resolution movie stacks using MotionCor2^21^ and binned twofold to a calibrated pixel size of 1.1 Å/pix. The defocus values were estimated by CTFFIND4^22^ from summed images without dose weighting. Other procedures of cryo-EM data processing were performed within RELION v3.1 or CryoSPARC v3 using the dose-weighted micrographs^23, 24^. For the cryo-EM dataset, a subset of ∼10,000 particles were picked by Laplacian-of-Gaussian (LoG) package in RELION without reference and subjected to reference-free 2D classification. Some of the resulting 2D class averages were low-pass filtered to 20 Å and used as references for automatic particle picking of the whole datasets in RELION, resulting in an initial set of 430,483 particles for reference-free 2D classification. 108,100 particles were selected from fine 2D classes for the initial 3D classification, using a 60 Å low-pass filtered initial model from cryoSPARC ab-initio reconstruction. After several rounds of 2D and 3D classification, 14,846 particles were 3D auto-refined, CTF refined, Bayesian polished, and post-processed, yielding a reconstruction of spike trimer in RBD-up conformation at 3.9 Å resolution. All reported resolutions are based on the gold-standard Fourier shell correlation (FSC) = 0.143 criterion. The GSFSC curves were corrected for the effects of a soft mask with high-resolution noise substitution. All cryo-EM maps were sharpened by applying a negative B-factor estimated in RELION or CryoSPARC. All the visualization and evaluation of the 3D volume map were performed within UCSF Chimera^25^ or UCSF ChimeraX^26^, and the local resolution variations were calculated using RELION^23^.

The initial templates for model building used the SARS-CoV-2 Omicron variant S trimer structure (PDB ID 7TGW)^7^ for the prefusion conformation and the wild-type S2 postfusion trimer structure (PDB ID: 6XRA)^27^ for the postfusion conformation. Briefly, the initial templates were fit into the map using Chimera and Coot^28^, followed by a ten-cycle rigid body refinement using Phenix. Then, a combined manual refinement and real-space refinement were carried out for both prefusion state and postfusion state S structures in Coot and Phenix^29^. The real-space refinement strategies included minimization global, local grid search, and adp, with rotamer, Ramachandran, and secondary structure restraints adapted from 7TGW and 6XRA. Statistics of map reconstruction and model refinement are listed in Supplementary Table S1. The final models were evaluated using MolProbity^30^. Map and model representations in the figures were prepared with UCSF Chimera and UCSF ChimeraX.

### Animal vaccination and serum collection

#### Mice

For mouse vaccination, groups of 6- to 8-week-old female BALB/c mice or female K18-hACE2 Transgenic Mice were intramuscularly immunized with LNP vaccine candidates or a placebo in 50 μL, and 3 weeks later, a second dose was administered to boost the immune responses.

#### Hamsters

6- to 8-week-old female hamsters were immunized intramuscularly with 150 μL of LNP vaccine candidates or placebo at week 0. At week 3, all hamsters received a boost vaccination.

#### Macaques

For the vaccination of rhesus macaques, three groups of 6∼8-year-old male or female rhesus macaques were immunized with LNP (n = 4/group, 2 male and 2 female), or PBS (n = 4, 2 male and 2 female) no more than 1mL (volume adjusted according to injected dose) via intramuscular injection twice at a three-week interval.

The sera of immunized mice, hamsters and rhesus macaques were collected and inactivated at 56°C for 0.5h to detect the SARS-CoV-2-specific IgG and neutralizing antibodies as described below.

### ELISA for SARS-CoV-2 S-specific IgG

SARS-CoV-2 S-specific antibody responses in immunized sera were determined by enzyme-linked immunosorbent assay (ELISA) assay, as previously described. Briefly, 96-well plates were coated with 50 μL of coating buffer containing 100 ng/well recombinant SARS-CoV-2 spike or RBD antigens (Sino Biological, Arco) at 4°C overnight. Plates were blocked with 2% bovine serum albumin solution in PBST at room temperature for 1 hour. Immunized mice sera were diluted 100-fold as the initial concentration, and then a 5-fold serial dilution of a total of 11 gradients in PBS buffer. PBST washed plates were incubated with serially diluted sera at room temperature for 2 hours. For determination of S-specific antibody response, plates were incubated with goat anti-mouse IgG HRP (for mouse sera, Proteintech Cat: SA00001-1) or goat anti-Syrian hamster IgG HRP (for hamster sera, abcam Cat: ab6892) or goat anti-monkey IgG HRP (for NHP sera, Invitrogen Cat: PA1-84631) at 37°C for 1 hour and then substrate tetramethylbenzidine (TMB) solution (Invitrogen) was used to develop. The color reaction was quenched with 1N sulfuric acid for about 10 minutes, and the optical density was measured at a wavelength 450 nm by Synergy H1 microplate reader (BioTek).

### Pseudovirus-based neutralization assay

The pseudovirus-based neutralization assay was performed at Genescript. Briefly, serum samples collected from immunized animals were serially diluted with the cell culture medium. The diluted serums were mixed with a pseudovirus suspension in 96-well plates at a ratio of 1:1, followed by 1 hours incubation at RT. Opti-HEK293/ACE2 cells were then added to the serum-virus mixture, and the plates were incubated at 37°C in a 5% CO_2_ incubator. 48 hr later, the luciferase activity, reflecting the degree of SARS-CoV-2 pseudovirus transfection, was measured using the Luciferase Assay kit. The NT_50_ was defined as the fold-dilution, which emitted an exceeding 50% inhibition of pseudovirus infection compared to the control group.

### SARS-CoV-2 neutralization assay

#### Cytopathic effect (CPE)

Serum samples collected from immunized animals were inactivated at 56°C for 30min and serially diluted with cell culture medium in two-fold steps. The diluted serums were mixed with a virus suspension of 100 TCID_50_ in 96-well plates at a ratio of 1:1, followed by 1 hours of incubation at 37°C in a 5% CO_2_ incubator. Vero E6 cells were then added to the serum-virus mixture, and the plates were incubated for 4 days at 37°C in a 5% CO_2_ incubator. Cytopathic effect (CPE) of each well was recorded under microscopes, and the neutralizing titer was calculated by the dilution number of 50% protective condition.

#### Plaque reduction neutralization test (PRNT)

The virus neutralization test was performed in a 24-well plate. The serum samples from immunized animals were heat-inactivated at 56°C for 30 min. The serum samples were diluted at 1:30, 1:90, 1:270, 1:810, 1:2430 and 1:7290, and then an equal volume of virus stock was added and incubated at 37°C in a 5% CO_2_ incubator. After 1 hour of incubation, 100 μL mixtures were inoculated onto monolayer Vero cells in a 24- well plate for 1 hour with shaking every 15 minutes. The inocula were removed, and cells were incubated with DMEM supplemented with 2% FBS containing 0.9% methylcellulose for 4 days before fixation. The cells were then fixed with 8% formaldehyde for 1 hour. The formaldehyde solution was removed and the cells were washed with tap water, followed by crystal violet staining. The plaques were counted for calculating the titer.

### ELISpot assay

The mouse and hamster elispot analysis was performed ex vivo using PBMCs with commercially available Mouse IFN-γ ELISpot assay kit (Dakewe) and Hamster IFN-γ ELISpotPLUS kit (Mabtech), respectively. The T cell immune responses in Cynomolgus macaques were detected using PBMCs with commercially available Monkey IFN-γ ELISpot assay kit and a Monkey IL-4 ELISpot assay kit (Mabtech). A pool of 15-mer peptides that overlapped by 11 amino acids and covered the whole sequence of the SARS-CoV-2 spike protein (Genscript) was used for ex vivo stimulation of PBMCs for ELISpot assay, which was divided into S1 peptide pool and S2 peptide pool. For the mouse and hamster IFN-γ ELISpot assays, 2.0×10^5^ PBMCs were stimulated with a final concentration of 1 μg/mL for pooled peptides of the SARS-CoV-2 spike. For the IFN-γ and IL-4 ELISpot assays in macaques, 2.5×10^5^ PBMCs were stimulated with a final concentration of 1 μg/mL for each spike peptide. The test for each animal was performed in two repetitions. RPMI-1640 medium served as an unstimulating control, and phytohemagglutinin (PHA) or PMA+Ionomycin (Dakewe) was used as positive controls. After 18-24 hours of stimulation with spike peptide pools, the substrate was added to the plate. The spots were counted by AT-Spo^TM^ 2100. The background (RPMI-1640 medium treated group) was subtracted and normalized to SFC/10^6^ PBMCs.

### Animal challenge experiments

#### BALB/C Mice

The mouse model for the SARS-CoV-2 B.1.351 variant challenge has previously been characterized. BALB/c mice immunized with RQ3013 (2 µg and 5 µg) and physiological saline were challenged with 5×10^4^ FFU SARS-CoV-2 B.1.351 variant via the intranasal at 2 weeks after the boost. The body weights of the mice were recorded daily. At 5 dpi, the immunized mice were sacrificed, and their lung tissues were collected to measure the viral RNA load and used for the histopathological assay.

#### K18-hACE2 Transgenic Mice

At 5 weeks after the boost, all the immunized K18-hACE2 Transgenic Mice (2 µg and 5 µg of RQ3013 and placebo) were challenged with 1.0×10^3^ PFU of B.1.351 SARS-CoV-2 virus via the intranasal routes. The body weights of the mice were recorded daily. At 4 dpi, all mice were sacrificed to collect specimens for further experiments.

#### Macaques

Rhesus macaques (6-8 years old) were divided into three groups and injected intramuscularly with high dose (100 μg/dose), low dose (30 μg/dose) vaccine, and physiological saline respectively. All grouped animals were immunized twice (days 0 and 21) before being challenged with 10^6^ TCID_50_/mL Wild-type SARS-CoV-2 virus by intratracheal routes. Macaques were euthanized, and lung tissues were collected at 7 dpi for RT-PCR and histopathological assays. The oropharyngeal, nasal, anal swabs, and blood samples were collected daily post-virus challenge.

### RT-PCR

#### BALB/C Mice

The viral RNA load in the lung tissues of infected BALB/c mice was detected by quantitative RT-qPCR. Briefly, the lung tissues were collected and homogenized with stainless steel beads in TRIZOL (1 ml for each sample). The RNAs in tissues were then extracted and reverse transcribed by AceQ Universal U+ Probe Master Mix V2 (Vazyme Biotech Co., Ltd). SARS CoV-2 RNA quantification was performed by RT-qPCR targeting the N gene of SARS-CoV-2, using HiScript III RT supermix for qPCR (+gDNA wiper) kit (Vazyme Biotech Co., Ltd). We used primers for the N gene: forward primer (F): 5’- GGGGAACTTCTCCTGCTAGAAT-3’; reverse primer (R): 5’-CAGACATTTTGCTCTCAAGCTG-3’; fluorescent probe (P): 5′-FAM- TTGCTGCTGCTTGACAGATT-TAMRA-3′. The amplification was performed as follows: 37°C for 2 min, 95°C for 5 minutes followed by 45 cycles (95°C for 10 s, 60°C for 34 s) and converted to virus copy number according to the standard curve on the real-time fluorescence quantitative PCR system-7500 instrument.

#### K18-hACE2 Transgenic Mice

The trachea, lung and brain tissues of challenged K18-hACE2 Transgenic mice were sufficiently broken with a biosafety tissue homogenizer. The nucleic acids in the samples were extracted according to the method of lysing the samples with TRIzol. Reverse transcription was performed with a PrimeScript RT Reagent Kit with gDNA Eraser (Takara, Cat no. RR047Q), and qRT-PCR was performed on StepOne Plus Real-time PCR system (Applied Biosystem) with TB Green Premix (Takara Cat no.RR820A). A standard curve was generated by the determination of copy numbers from serially dilutions (103 - 109 copies) of the plasmid. The primers used for quantitative PCR were RBD-qF: 5’- CAATGGTTTAACAGGCACAGG- 3’ and RBD-qR: 5’-CTCAAGTGTCTGTGGATCACG-3’. PCR amplification was performed as follows: 95°C for 5 min followed by 40 cycles consisting of 95°C for 15 s, 54°C for 15 s, and 72°C for 30 s. The dose-response curves were plotted from viral RNA copies versus the drug concentrations using GraphPad Prism 8.0 software.

#### Macaques

Viral RNA in the rhesus samples was quantified by one-step real-time quantitative RT-PCR. According to the manufacturer’s instructions, the swab and blood samples were used to extract viral RNA by using the QIAamp Viral RNA Mini Kit (Qiagen). Tissues were homogenized in DMEM (1:10, W/V), the homogenate was clarified by low-speed centrifugation at 4500 × g for 30 min at 4°C, and the supernatant was immediately used for RNA extraction. RNA was eluted in 50 μL of elution buffer and used as the template for RTPCR. The primer pairs targeting the S gene were used as follows: RBD-qF1: 5′- CAATGGTTTAACAGGCACAGG-3′; RBDqR1: 5′-CTCAAGTGTCTGTGGATCACG-3′. Two microliters of RNA were used to verify the RNA quantity by HiScript® II One Step qRTPCR SYBR® Green Kit (Vazyme Biotech Co., Ltd) according to the manufacturer’s instructions. The amplification was performed as follows: 50°C for 3 min, 95°C for 30 s followed by 40 cycles (95°C for 10 s, 60°C for 30 s), with a default melting curve step in an ABI StepOne system.

### Histopathology and immunohistochemistry

#### BALB/C Mice

To study pathological lung damage, lung tissue samples of BALB/c mice were fixed with 4% paraformaldehyde and embedded, sectioned, stained, and sealed according to the testing SOP procedures. Tissue slices were observed in detail at different magnifications. Pathological changes such as congestion, hemorrhage, edema, degeneration, necrosis, hyperplasia, fibrosis, organization were marked. The slide images were collected using Pannoramic DESK and analyzed with Caseviewer C. V 2.4. Typical lesions are imaged and marked with arrows.

#### K18-hACE2 Transgenic Mice

Animal necropsies were performed according to a standard protocol. The collected lung tissue was fixed in 10% neutral formalin buffer for more than 3 days and removed from the laboratory (in accordance with the regulations and requirements of the removal of infectious experimental materials inactivated by the P3 experiment), paraffin-embedded and sections were stained with hematoxylin and eosin and observed under a light microscope.

#### Macaques

To study the pathogenesis and pathological damage of the respiratory tract, animals in each group were euthanized at 7 dpi and six lung lobes were collected for pathological, virological, and immunological analysis. The dissection of experimental animals was carried out according to the standard operation of experimental animals. The collected organ and tissue samples were fixed in 10% neutral buffered formalin and processed routinely into paraffin blocks. Tissue sections were prepared, and slides were stained with hematoxylin and eosin (H&E) before examination by light microscopy.

### Vaccine safety evaluation

The safety of RQ3013 was evaluated in cynomolgus macaques. Five groups of monkeys (5 female and 5 male monkeys/group) were immunized with high dose (240 μg /dose), low dose (60 μg /dose) vaccine, or high dose, low dose empty LNP only and physiological saline individually for three times at days 0, 14 and 28. Datasets of many safety-related parameters were collected during and after immunization, including clinical observation, body weight, and body temperature. Analysis of lymphocyte subset percent (CD3^+^, CD4^+,^ and CD8^+^), key cytokines (TNF-α, IFN-γ, IL-2, IL-4, IL-5, and IL-6), and biochemical blood tests are also performed in collected blood samples. 60% of monkeys were euthanized at day 31 post-immunization, and the left 40% were euthanized on day 42. Organs of lung, brain, heart, liver, spleen, and kidney were collected for pathologic analysis.

### Statistical Analysis

All statistics data were performed and graphed using GraphPad Prism8.0. The EC50 values were calculated by non-linear regression. Statistical analyses were carried out by Student’s t-test when two groups were analyzed and by ANOVA when more than two groups were analyzed.

## Acknowledgments

This work was supported by the National Key Research and Development Program of China (2021YFC0865700 to J. Lin, 2021YFE0201900 to C. Shan, 2021YFC2301700 to M. Wang), China Postdoctoral Science Foundation (2020T130017ZX to J. Lu), the Starry Night Science Fund at Shanghai Institute for Advanced Study of Zhejiang University (SN-ZJU-SIAS-009 to J.Lin), the STS regional key project (KFJ-STS-QYZD-2021-12-001 to Z. Yuan and C. Shan) from Chinese Academy of Sciences, and Hubei Science and Technology Major Project (2021ACB0004 to C. Shan). We are particularly grateful to the staff at the Wuhan National Biosafety Laboratory of the Chinese Academy of Sciences. We thank the Center for Instrumental Analysis and Metrology and BSL-3 laboratory at the Wuhan Institute of Virology. We also thank the Cryo-Electron Microscopy Facilities at the Multiscale Research Institute of Complex Systems (300 kV) and the School of Life Sciences (200 kV) at Fudan University.

## Author Contributions

J.LIN and J.LU conceived and designed the experiments. S.T. performed most of the sera assays. J.Z., X.G., B.D., Y.LIU, L.L., Y.C., and Y.LI participated in the production and quality control of mRNA-LNPs. Z.W., G.L., and Z.Y. determined the cryo-EM structures. X.H., Y.F.Y., X.Z., J.R., G.G., Y.P., H.L., Z.Y., and C.S. performed experiments related to rhesus macaques. Y.F.L., J.LIU., Q.W., H.H., and M.W. performed experiments related to K18-hACE2 mice. H.Z., Y.Y., Y.X., W.Y., L.F., J.LU, and J.LIN analyzed the data. J.LIN, W.Y., H.Z., and S.T. wrote the manuscript. All authors provided the final approval of the manuscript.

## Data Availability

The atomic coordinates and cryo-EM maps of the S immunogen have been deposited in the Protein Data bank (PDB) and Electron Microscopy Data Bank (EMDB). Accession numbers are 7L7F and EMD-33346 for the prefusion state, and 7XOG and EMD-33347 for the postfusion state.

