## Supplemental Information for "Preclinical evaluation of RQ3013, a broad-spectrum mRNA vaccine against SARS-CoV-2 variants"

### Supplementary information Fig. S1

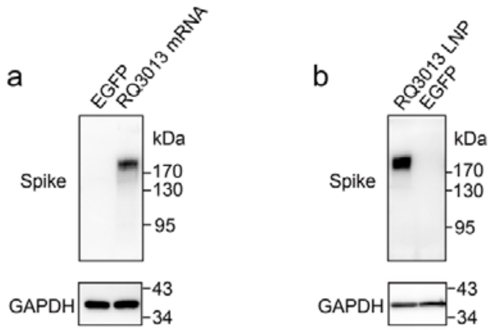


**Fig. S1 Expression of the protein antigen in HEK 293A cells transfected with RQ3013 mRNA or LNP.** RQ3013 mRNA (**a**) and LNP (**b**) were used to transfect HEK 293A cells, and protein expression was detected by western blot. EGFP mRNA was used as a negative control.

### Supplementary information Fig. S2

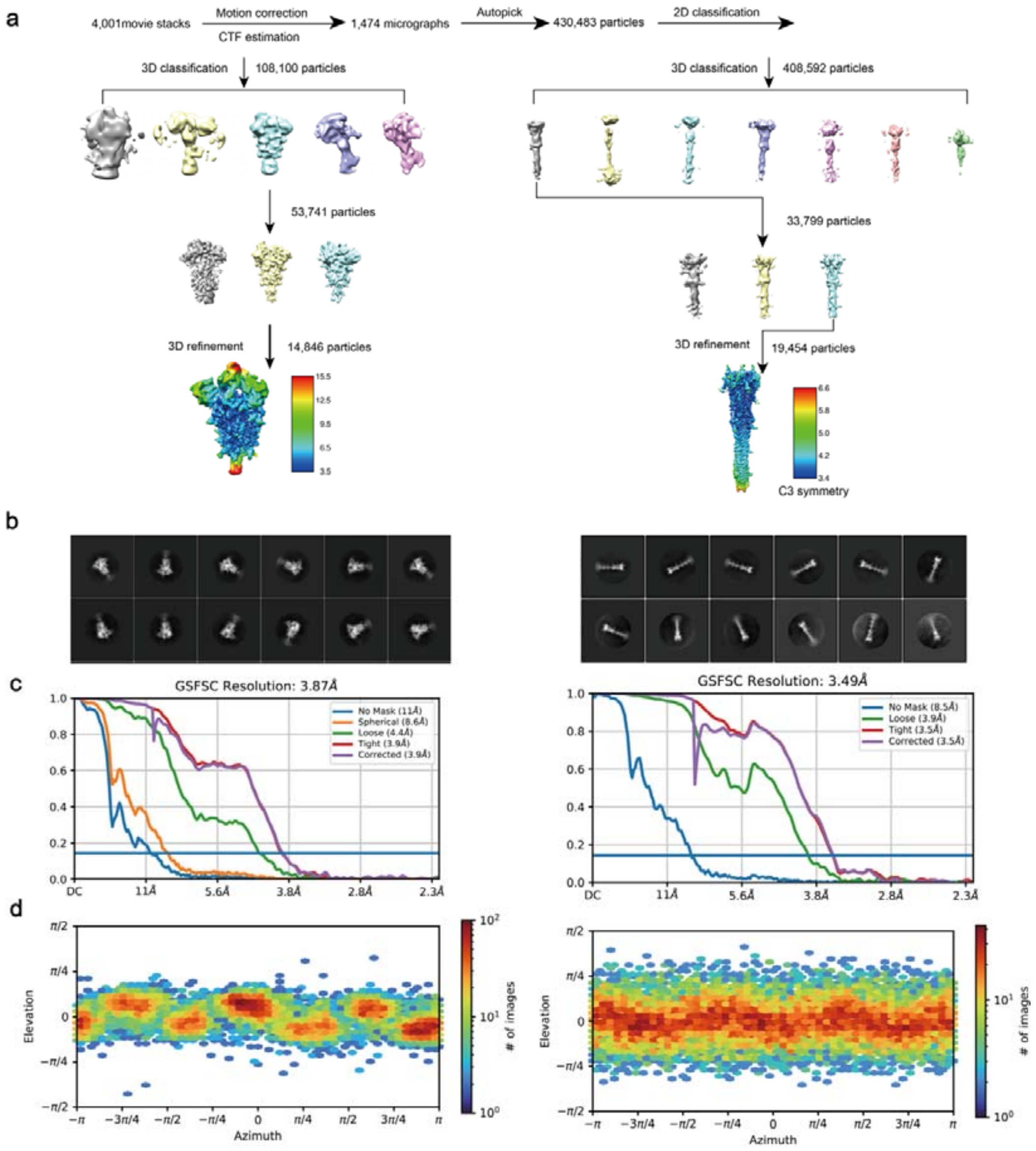


**Fig. S2 Cryo-EM data collection, image processing, and 3D reconstruction. a** The flowchart of the cryo-EM image processing and 3D reconstruction for the RQ3013 immunogen in prefusion and postfusion conformations. **b** Representative cryo-EM 2D classification of the S trimer in prefusion and postfusion conformations. All the data were collected on a K3 direct electron detector. (**c, d**) The GSFSC curve and angular distribution for the 3D reconstruction of the cryo-EM map of the S trimer in prefusion and postfusion conformations.

### Supplementary information Fig. S3

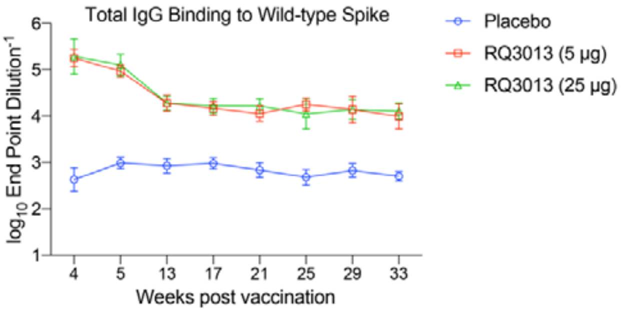


**Fig. S3 Long-term monitoring of immunogenicity of RQ3013 in hamsters.** Hamsters were immunized two times on days 0 and 21 through the intramuscular route with a low dose (5 μg per dose) or high dose (25 μg per dose) of RQ3013 or PBS. The antibody response in sera from weeks 4 to 33 following the prime vaccination was analyzed by ELISA using the wild-type S protein (n >= 7). Values are GMT mean ± SD.

### Supplementary information Fig. S4

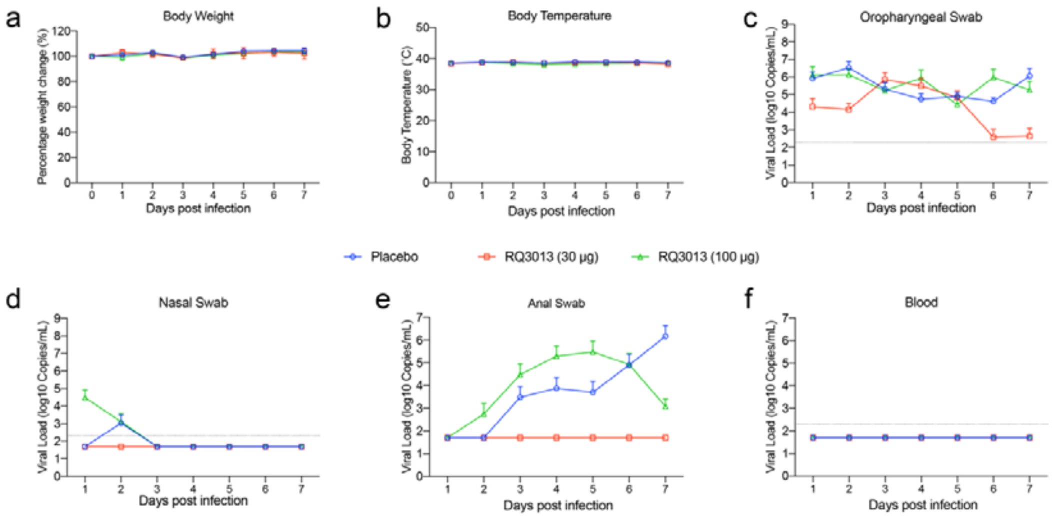


**Fig. S4 Monitoring of body weight, body temperature, and viral RNAs in swabs and blood of rhesus macaques challenged with live virus. a** Body weight changes of macaques after infection with live wild-type SARS-CoV-2. **b** Body temperature changes. **c-f** Viral load in oropharyngeal, nasal, and anal swabs, and in blood. The black dashed line indicates the assay's detection limit (200 Copies/mL). Any measurement below the detection limit was assigned a value of half the limit of detection for plotting and statistical purposes. Values are mean ± SD.

### Supplementary information Fig. S5

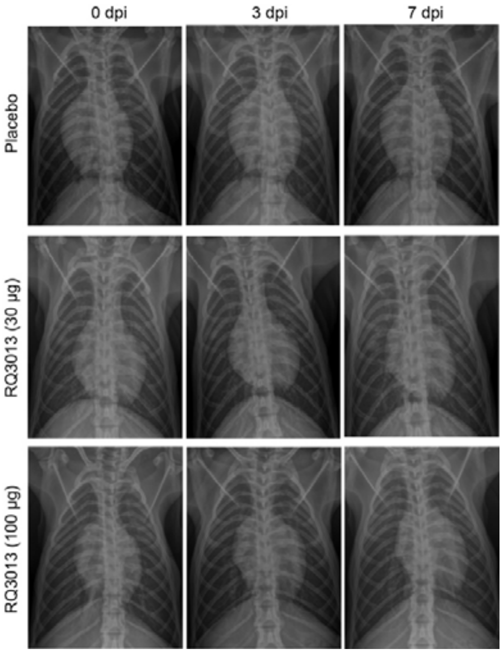


**Fig. S5 Chest radiographs of rhesus macaques.** X-ray examinations were performed at 0, 3, and 7 dpi by HF100Ha (MIKASA, Japan) for all the available RMs.

### Supplementary information Fig. S6

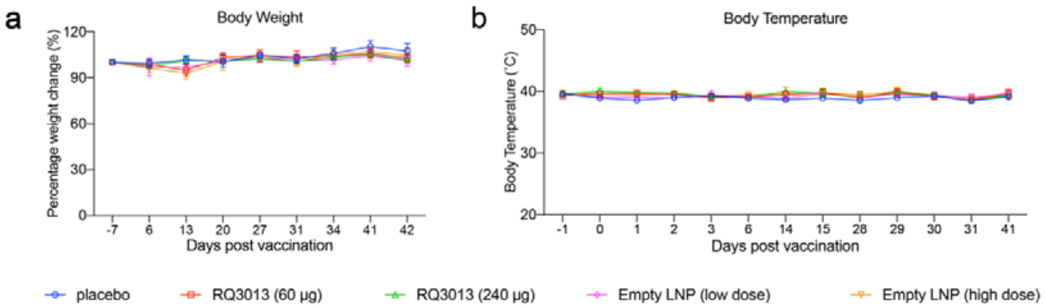


**Fig. S6 Body weight and temperature changes in immunized cynomolgus macaques.** Macaques (n = 10 per group) were vaccinated three times on days 0, 14, and 28 through the intramuscular route with PBS (Placebo) or low dose (60 μg) or high dose (240 μg) of RQ3013 or low and high doses of empty LNP. **a** Body weight. **b** Body temperature. Values are mean ± SD.

### Supplementary information Fig. S7

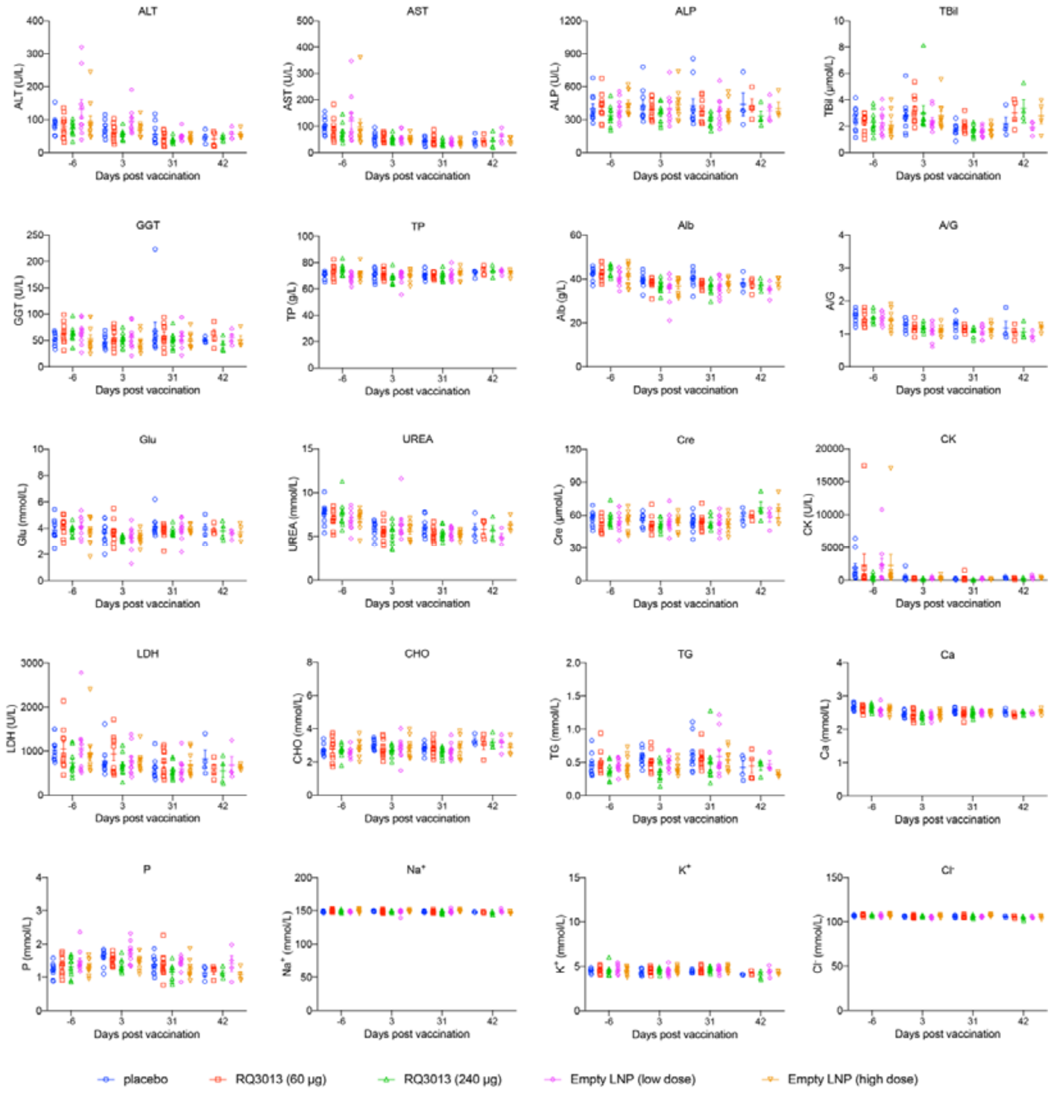


**Fig. S7 Hematological indices in immunized cynomolgus macaques.** Macaques (n = 10 per group) were vaccinated three times on days 0, 14, and 28 through the intramuscular route with PBS (Placebo) or low dose (60 μg) or high dose (240 μg) of RQ3013 or low and high doses of empty LNP. The selective hematological indices were measured at different time points. ALT (Alanine aminotransferase), AST (Aspartate aminotransferase), ALP (Alkaline phosphatase), TBil (Total bilirubin), GGT (γ-glutamyltranspeptidase), TP (Total protein), Alb (Albumin), A/G (Albumin/globulin ratio), Glu (Glucose), UREA (Blood urea), Cre (Creatinine), CK (Creatine kinase), LDH (Lactate dehydrogenase), CHO (Total cholesterol), TG (Triglycerides), Ca (Calcium), P (Phosphorus), $Na^+$ (Sodium ion), $K^+$ (Potassium ion), $Cl^-$ (Chloride ion). Values are mean ± SEM.

### Supplementary information Fig. S8

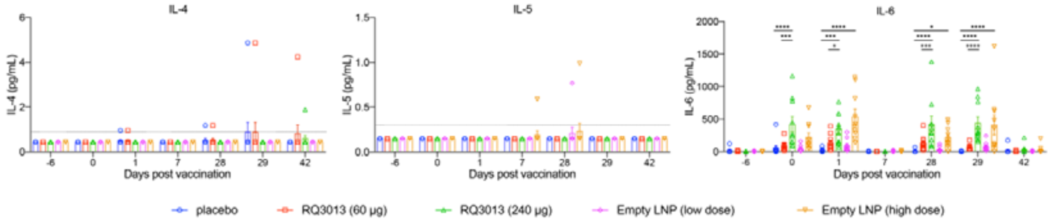


**Fig. S8 Key cytokine levels in immunized cynomolgus macaques.** Macaques (n = 10 per group) were immunized three times on days 0, 14, and 28 through the intramuscular route with PBS (Placebo) or low dose (60 μg) or high dose (240 μg) of RQ3013 or low and high doses of empty LNP. Key cytokines of IL4, IL5, and IL6 were measured at different time points. The black dashed line indicates the detection limit of the assay (IL-4, 0.9 pg/mL; IL-5, 0.3 pg/mL; IL-6, 0.1 pg/mL). Any measurement below the detection limit was assigned a value of half the limit of detection for plotting and statistical purposes. Values are mean ± SEM.

### Supplementary information Fig. S9

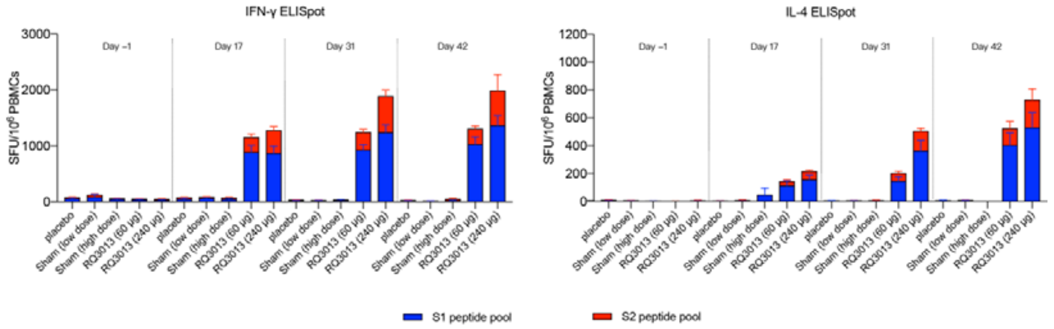


**Fig. S9 Cellular immune responses of RQ3013 in cynomolgus macaques.** Macaques (n = 10 per group) were immunized three times on days 0, 14, and 28 through the intramuscular route with PBS (Placebo) or low dose (60 μg) or high dose (240 μg) of RQ3013 or low and high doses of empty LNP. PBMCs obtained on day -1 (pre-prime), 17 (3 days after the second vaccination), 31 (3 days after the third vaccination), and on day 42 (14 days after the third vaccination) were stimulated with overlapping peptide pools of S1 and S2 from the RQ3013 antigen. The IFNγ or IL-4 ELISpot assays were then performed. SFU, spot-forming units. Values are mean ± SEM.

### Supplementary information Fig. S10

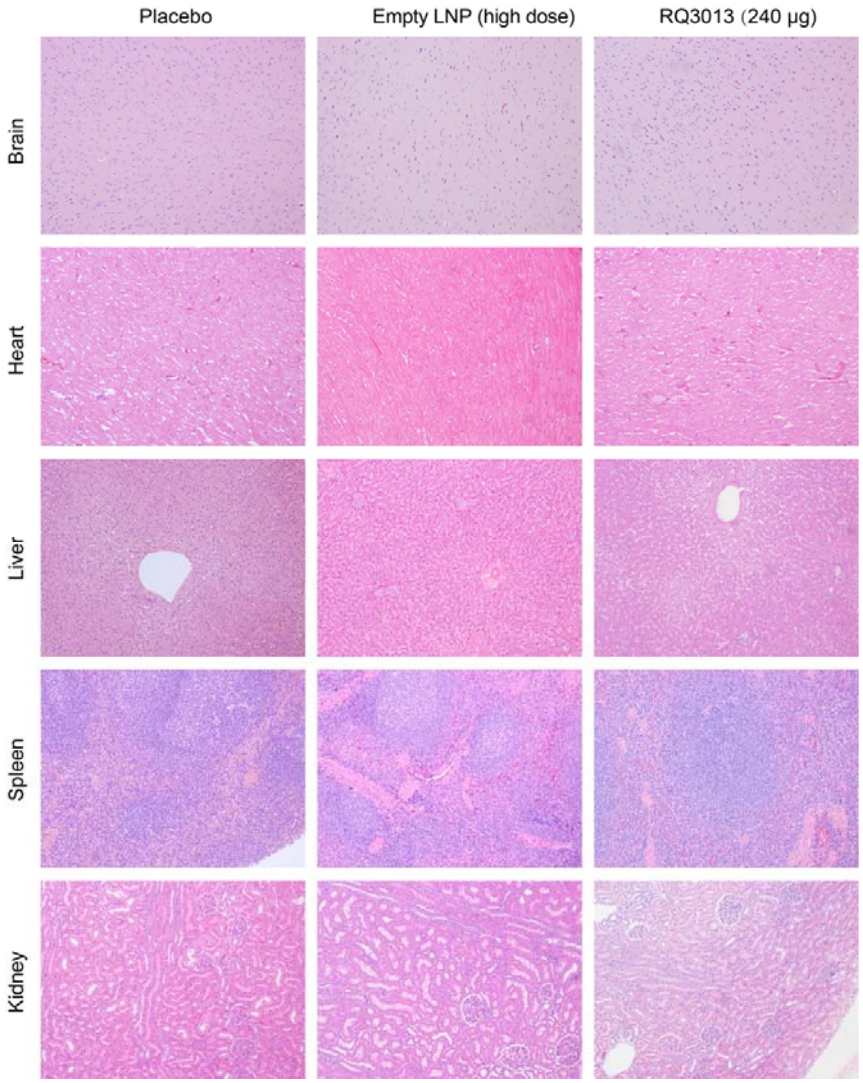


**Fig. S10 Histopathological evaluations of RQ3013 safety in nonhuman primates.** Cynomolgus macaques (n = 10 per group) were immunized three times on days 0, 14, and 28 through the intramuscular route with PBS (Placebo) or low dose (60 μg) or high dose (240 μg) of RQ3013 or low and high doses of empty LNP. The brain, heart, liver, spleen, and kidney tissues from three groups of aminals were collected on day 31 and stained with hematoxylin and eosin for histopathological evaluations.

### Supplementary information Table S1

Table S1 Refinement and model statistics of the S glycoprotein encoded by the Covid-19 mRNA vaccine candidate RQ3013

| **Data collection and processing** | prefusion | postfusion |
|---|---|---|
| PDB and EMDB | 7L7F and EMD-33346 | 7XOG and EMD-33347 |
| Magnification | 81,000x | 81,000x |
| Voltage (kV) | 300 | 300 |
| Electron exposure ($e^-$/pix/s) | ~ 20 | ~ 20 |
| Number of frames per movie | 40 | 40 |
| Defocus range (μm) | -1.5 to -2.5 | -1.5 to -2.5 |
| Pixel size (Å) | 1.1 | 1.1 |
| Symmetry imposed | C1 | C3 |
| Number of used micrographs (no.) | 1,474 | 1,474 |
| Total of extracted particles (no.) | 430,483 | 795,627 |
| Total of refined particles (no.) | 14,846 | 19,454 |
| Resolution Masked 0.143 FSC (Å) | 3.9 | 3.5 |
| **Refinement** | | |
| Map sharpening B-factor (Å$^2$) | -15.4 | -64.5 |
| Non-hydrogen atoms | 25,760 | 8971 |
| Protein residues | 3217 | 1068 |
| Lignads/Glycans | 45 | 57 |
| **r.m.s. deviations** | | |
| Bond lengths (Å) | 0.003 | 0.004 |
| Bond angles (°) | 0.762 | 0.757 |
| **Validation** | | |
| MolProbity score | 2.06 | 2.12 |
| All-atom clashscore | 11.88 | 11.30 |
| Rotamers outliers (%) | 0.43 | 0.64 |
| Cβ outliers (%) | 0.03 | 0.00 |
| CaBLAM outliers (%) | 3.98 | 3.74 |
| **B-factors (min/max/mean)** | | |
| Protein | 47.2/395.7/181.1 | 0.05/142.82/13.65 |
| Ligands/Glycans | 94.8/343.1/170.3 | 16.41/213.85/149.39 |
| **Overall correlation coefficients** | | |
| CC (mask) | 0.82 | 0.79 |
| CC (box) | 0.87 | 0.80 |
| CC (peaks) | 0.75 | 0.80 |
| CC (volume) | 0.80 | 0.62 |
| **Ramachandran plot statistics** | | |
| Favored (%) | 92.48 | 90.06 |
| Allowed (%) | 7.49 | 9.66 |
| Disallowed (%) | 0.03 | 0.28 |
